# How does genome size affect the evolution of pollen tube growth rate, a haploid performance trait?

**DOI:** 10.1101/462663

**Authors:** John B. Reese, Joseph H. Williams

## Abstract

**Premise of the Study:** Male gametophytes of most seed plants deliver sperm to eggs via a pollen tube. Pollen tube growth rates (*PTGRs*) of angiosperms are exceptionally rapid, a pattern attributed to more effective haploid selection under stronger pollen competition. Paradoxically, whole genome duplication (WGD) has been common in angiosperms but rare in gymnosperms. Pollen tube polyploidy should initially accelerate *PTGR* because increased heterozygosity and gene dosage should increase metabolic rates, however polyploidy should also independently increase tube cell size, causing more work which should decelerate growth. We asked how genome size changes have affected the evolution of seed plant *PTGRs*.

**Methods:** We assembled a phylogenetic tree of 451 species with known *PTGRs*. We then used comparative phylogenetic methods to detect effects of neo-polyploidy (within-genus origins), DNA content, and WGD history on *PTGR*, and correlated evolution of *PTGR* and DNA content.

**Key Results:** Gymnosperms had significantly higher DNA content and slower *PTGR* optima than angiosperms, and their *PTGR* and DNA content were negatively correlated. For angiosperms, 89% of model weight favored Ornstein-Uhlenbeck models with a faster *PTGR* optimum for neo-polyploids, but *PTGR* and DNA content were not correlated. In comparisons of within-genus and intraspecific-cytotype pairs, *PTGRs* of neo-polyploids ≤ paleo-polyploids.

**Conclusions:** Genome size increases should negatively affect *PTGR* when genetic consequences of WGDs are minimized, as found in intra-specific autopolyploids (low heterosis) and gymnosperms (few WGDs). But in angiosperms, the higher *PTGR* optimum of neo-polyploids and non-negative *PTGR*-DNA content correlation suggest that recurrent WGDs have caused substantial *PTGR* evolution in a non-haploid state.

## INTRODUCTION

In seed plants, the male gametophyte is a highly-reduced, haploid organism that develops within the pollen grain and completes its life cycle after pollination by growing a pollen tube that invades female reproductive tissues. The pollen tube functions to attach the male gametophyte and to absorb nutrients from female tissues, and in most seed plants (conifers, Gnetales, and angiosperms), it has the novel function of transporting the sperm cells to the egg-bearing female gametophyte (siphonogamy) (Friedman, 1993). Pollen tube growth rate (*PTGR*) is a central aspect of male gametophyte performance that can evolve due to changes in the time between pollination and fertilization, and due to changes in the intensity of pollen tube competition. Strikingly, angiosperms are known to have much shorter reproductive cycles (Williams and Reese, 2019), much higher potential for pollen competition (Mulcahy, 1979), and orders of magnitude faster *PTGRs* (Williams, 2012) relative to gymnosperms. The pattern of exceptionally fast angiosperm *PTGRs* is thought to have evolved rapidly via haploid selection on pollen-expressed genes (Mulcahy, 1979; Arunkumar et al., 2013; Otto et al., 2015), which constitute a large portion of the genome (Tanksley et al., 1981; Rutley and Twell, 2015; Hafidh et al., 2016).

If the dramatic and rapid acceleration of *PTGR*s in angiosperms has been driven by haploid selection on pollen performance genes, then one might expect polyploidy to be rare in angiosperms. Evolution above the haploid level is expected to reduce the efficiency of selection on pollen (Otto et al., 2015). Yet, the opposite is true – ancient whole genome duplications (WGDs), recent polyploids, and speciation by polyploidy have been especially common in angiosperms, whereas in gymnosperms genome size has evolved largely by other processes (Wood et al., 2009; Mayrose et al., 2011; Leitch & Leitch, 2012, 2013; Landis et al., 2018). In fact, large changes genome size can have a number of immediate effects on *PTGR*. First, *PTGR* might be faster in a neo-diploid pollen tube since increases in gene number cause: 1) heterosis, due to sheltering of deleterious pollen-expressed alleles and/or new allelic interactions upon loss of haploidy (Lande and Schemske, 1985; Husband and Schemske, 1997; Comai, 2005; Birchler et al. 2010; Husband, 2016), and 2) gene dosage effects, due to increased capacity for protein synthesis and hence the possibility for higher metabolic rates (Stebbins, 1974; Comai, 2005; Conant and Wolfe, 2008). On the other hand, substantial increases in DNA content (whether by WGD or other processes) are known to increase nuclear size, cell size, and the duration of the cell cycle, independent of the effects of genes (Bennett, 1971, 1972; Cavalier-Smith, 1978; Price 1988; Cavalier-Smith, 2005). The phenotypic effects of increased bulk DNA, hereafter referred to as “nucleotypic” effects (Bennett, 1971; Snodgrass et al, 2016; Doyle & Coate, 2019), cause more work for the growing pollen tube cell and should therefore negatively affect *PTGR*, counteracting the positive “genotypic” effects of heterozygosity and gene dosage.

As shown in Figure 1, if genome size expansion occurs without increasing the number of genes, then nucleotypic effects will predominate and slower *PTGRs* should evolve. But if genome size increase occurs by WGD, then altered gene expression patterns (due to dosage and heterozygosity effects) should counteract nucleotypic effects in the stabilized neo-polyploid (Fig. 1). In the latter case, the balance of nucleotypic and genotypic effects varies depending on the magnitude of potential heterosis, which depends directly on the amount of standing genetic variation (Birchler et al. 2010). In general, at inception tetraploid sporophytes are expected to have higher heterozygosity than their diploid progenitors, irrespective of mode of polyploidization (auto-to allo-polyploidy) or mating system (Lande and Schemske 1985; Soltis and Soltis 2000). Thus, at inception, autopolyploids that arise from outcrossing progenitors and allopolyploids will have a higher potential for heterosis, relative to autopolyploids that arose from selfing ancestors (Fig. 1).

**Figure 1.**
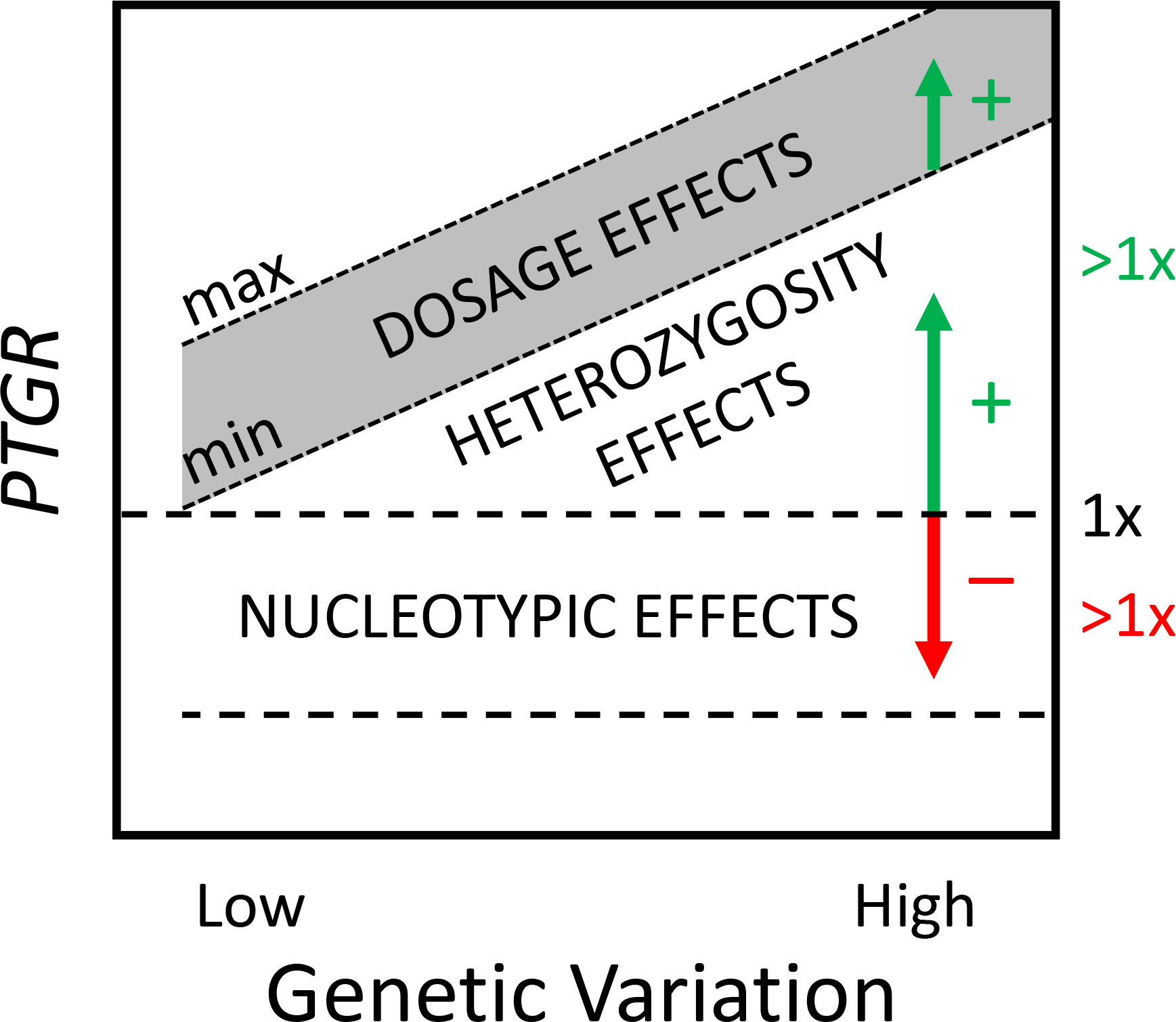
Predicted initial effects of large increases in genome size on pollen tube growth rate (*PTGR*). The dashed line indicates an ancestral haploid (1x) *PTGR*. Upon transition to a larger (> 1x) genome size, nucleotypic effects should act to decrease *PTGR* regardless of mechanism of change. Genotypic effects are only present after WGD or large-scale gene duplications and are predicted to increase *PTGR* via increased gene dosage and heterozygosity. The magnitude of heterosis due to initial increase in heterozygosity is expected to scale with genetic variation in the descendent taxon. The ancestral haploid *PTGR* can only be conserved when genotypic and nucleotypic effects perfectly offset each other.

After the initial effects of WGD, genotypic effects continue to evolve under both stabilizing and directional selection on *PTGR*, mediated by shifts in mating system and phenomena such as genome downsizing, biased gene retentions, recombination, and ultimately the return to disomic inheritance (Conant and Wolfe, 2008; De Smet et al. 2013; Conant et al., 2014; Freeling et al., 2015; Dodsworth et al., 2016; Panchy et al. 2016; Wendel et al., 2018). Nucleotypic effects by definition can only evolve by changes in genome size, which after WGD tend to be biased to small losses relative to the size of the WGD (Dodsworth et al. 2016). Hence, with time, genotypic effects are predicted to overwhelm nucleotypic effects, irrespective of initial effects and the direction of *PTGR* evolution.

In this study, we used model-based comparative phylogenetic analyses to determine if polyploidy, DNA C-value, and WGD history have affected the evolution of *PTGRs* in seed plants, and whether genome size effects have occurred predominantly during polyploid periods of history or during subsequent periods of more or less diploid evolution. Because all seed plants have at least one WGD in their history, we defined neo-polyploids as having a higher chromosome multiple than the base chromosome number of their genus, and paleo-polyploids (hereafter, “diploids”) as having similar chromosome number as the genus base number (as in Wood et al. 2009; Mayrose et al. 2011). This allowed us to determine if, 1) neo-polyploids have faster *PTGRs* than diploids, as predicted if WGDs generally produce strong initial genotypic effects that persist in the polyploid condition, or 2) neo-polyploids have slower *PTGRs* than diploids, as predicted if nucleotypic effects initially outweigh genotypic effects and if fast *PTGRs* generally evolve after diploidization (eg. in paleopolyploids). We also predict an underlying negative correlation between *PTGR* and genome size due to nucleotypic effects, which should be most apparent in intraspecific neo-polyploids and in lineages with little history of WGD.

## MATERIALS AND METHODS

### Tree Construction and Dating

GenBank accessions for 16 gene regions (*rbcL, matK, trnL-F, 18s_rDNA, atpB, ndhF, adh, trnL, rpl32, trnT-L, psbA-trnH, rpl32-trnL, ITS, 5.8s_rRNA, rps16*, and *26s_rDNA*) for 451 seed plant species with pollen tube growth rate data were retrieved, cleaned, and assembled into a multiple gene alignment (length – 9263 base pairs, 16 partitions, 69.6% missing data) using PHLAWD and phyutility (Smith and Donoghue, 2008; Smith and Dunn, 2008). Tree inference was performed using maximum likelihood in RAxML version 8 (Stamatakis, 2014) on CIPRES. A pruned version of the seed plant tree from Magallón et al. (2015) was used as a guide tree to enforce topology of major clades. The resulting maximum likelihood estimate of the tree was rooted and ultrametricized using the *ape* (Paradis et al., 2004) and *geiger* packages in R (Harmon et al., 2008). Time-calibration was performed with the Congruification method (Eastman et al., 2013), using the Magallón et al. (2015) phylogeny as the reference tree.

### Data collection and character scoring

Data on *PTGRs* were taken from Williams (2012) and more recent literature (cited in Appendix S1; see the Supplementary Data with this article). The *PTGR* value used for each species represents an estimate of maximum sustained growth rate, which is consistent with other comparative analyses of physiological traits, and with the fact that researchers almost always measure *PTGRs* from the longest pollen tube(s). Thus, *PTGR* values for each species represent an average of maximum in vivo growth rates, or if there was more than one report for a species the average of those values (as in Williams, 2012). *PTGRs* were taken exclusively from within-ploidy level crosses (i.e., never from inter-ploid crosses), in keeping with our overall goal of finding mechanisms underlying the pattern of *PTGR* evolution within stabilized polyploids.

DNA content was analyzed using 1C-value: the amount of nuclear DNA in the unreplicated gametic nucleus, irrespective of ploidy level (Swift, 1950; Bennett and Leitch, 2012). As we were primarily interested in the nucleotypic effects of bulk DNA amount, we use the terms C-value, DNA content, and genome size interchangeably throughout. C-value data was collected from the Kew Royal Botanic Gardens Plant C-value Database (Bennett and Leitch, 2012). Chromosome counts were obtained from the Index to Plant Chromosome Numbers (IPCN). To examine the effect of recent polyploidy (defined as occurring at or within the genus level; Wood et al., 2009; Mayrose et al., 2011) on *PTGR*, we scored taxa as “neo-polyploid” if their chromosome counts were ≥ 1.5 times that of their generic 1x base count (from Wood et al., 2009) and “diploid” (paleo-polyploid) if < 1.5 times that value (*N* = 206 angiosperms, 23 gymnosperms). To examine the effect of ancient (deeper than genus-level) duplication events on *PTGR*, the number of WGDs in each genus-to-root lineage was counted for each angiosperm (found in Appendix S1 of Landis et al. 2018) and gymnosperm (Li et al. 2015).

### Phylogenetic Comparative Analyses

To visualize changes in DNA content and *PTGR* along tree branches and to generate estimates of node states, ancestral state reconstructions were performed and plotted using the contMap function in *phytools* (Felsenstein, 1985; Revell, 2012)(Appendix S2). Given many known biological differences between gymnosperms and angiosperms in pollen tube growth (Friedman, 1993; Williams, 2008) and in mechanisms of genome size change (see Discussion) (Ohri and Khoshoo, 1986; Leitch et al., 1998), all comparative analyses were performed on gymnosperms only, angiosperms only, and the full dataset (all spermatophytes). C-value and *PTGR* were log_10_-transformed for all analyses.

Model-based analyses were used to examine patterns of *PTGR* and C-value evolution separately. The *OUwie* function was implemented in R (Beaulieu and O’Meara, 2014), and the following models were tested: single- and multi-rate Brownian motion (BM1, BMS, respectively), single-regime Ornstein-Uhlenbeck (OU1), and multi-regime OU models with either one global ***α*** and ***σ^2^*** estimate (OUM), one ***α*** and multiple ***σ^2^*** (OUMV), or multiple ***α*** and one ***σ^2^*** (OUMA). In all models, ***σ^2^*** represents the rate of random evolution and ***α***,the strength of attraction to an optimum, θ. The single- and multiple-regime models were compared to test whether or not, 1) angiosperms and gymnosperms evolve around different *PTGR* or C-value optima, and 2) diploids and neo-polyploids (within all three groups) evolve around different *PTGR* or C-value optima. For all analyses, AICc values were used to calculate model weights and the weighted average of parameter values was then calculated using all models that contributed > 1% of the model weight (Burnham & Anderson, 2002). Unless otherwise noted, all measures of uncertainty around parameter estimates are standard errors.

Since each *PTGR* value represents a species mean obtained from multiple measurements, we attempted to incorporate error into phylogenetic comparative analyses. Since species means were log_10_-transformed for analysis, log_10_-transformed *SE*s are also required. As there is no reliable way to calculate the log_10_-transformed *SE* from the literature without the original data for each species, we took the following approach. First, we assumed all species had similar *SE*s in *PTGR*, and we applied an empirically-determined *SE* from an exemplar species to all. *Magnolia grandiflora* has an average *PTGR* of 828 ± 141 μm h^−1^ (*N =* 25 outcrosses), close to the angiosperm median of 587 μm h^−1^ (Williams, 2012 and this study) (Appendix S3). The standard deviation (*SD*) of log_10_-transformed data was calculated and divided by the mean of the log_10_-transformed data to acquire a coefficient of variation (*CV*) of 0.0237. We then multiplied the log_10_-transformed mean *PTGR* of each species by 0.0237 to provide an estimate of the log_10_ taxon-specific standard deviation. The standard deviation (*SD*) was used as a conservative estimate of error because sample sizes were generally not available for calculating *SE*. Secondly, we performed a sensitivity analysis by evaluating each evolutionary model in *OUwie* with *SDs* calculated from hypothetical global *CV*s of 0.00, 0.05, 0.10, 0.25, and 0.50 (Appendix S4).

The association between recent polyploidy and *PTGR* was also assessed among 10 diploid-polyploid near-relative pairs (appearing as sister taxa on the tree at the within-genus level). Only polyploid taxa with a single diploid sister on the tree were used. The *PTGRs* of 11 intraspecific diploid-autopolyploid pairs from the literature were also compared. A two-tailed binomial (sign) test was used to test significance in both.

The cumulative effect of ancient polyploid events was explored with phylogenetic generalized least squares (*PGLS*) regression using the *phylolm* package in R (Ho & Ane, 2014). The number of ancient duplication events in the history of each tip taxon (inferred from Landis et al., 2018) was used as the predictor variable with *PTGR* as the response variable.

The relationship between pollen tube growth rate and gametophytic DNA content was also assessed with *PGLS* regression. Gametophytic DNA content was used as the predictor variable and *PTGR* the response variable. BM (Grafen, 1989) and OU (Martins and Hansen, 1997) models were both used, in addition to Pagel’s lambda, kappa, and delta models (Pagel, 1997, 1999). To examine the effect of ploidy and the interaction between ploidy and C-value on *PTGR*, a phylogenetic ANCOVA was implemented with C-value as the covariate in *phylolm*.

Shifts among convergent *PTGR* and C-value optima were determined with a maximum likelihood approach to detect multiple optima within seed plants, using *SURFACE* in R (Ingram and Mahler, 2013). Using an OU model with a global α and σ^2^, a single-optimum model was subdivided into multiple-optima models in a stepwise fashion until adding another optimum decreased the model likelihood by ∆AIC > −2. Separate optima were then collapsed (i.e. two regimes were assigned the same optimum) in a pairwise fashion until further collapses decreased model likelihood. Shifts in *PTGR* and C-value optima that occurred at the same node, or within two nodes of each other, were identified manually. Nodes with *PTGR* or C-value regime shifts were also manually compared to the Landis et al. (2018) WGD map to see if a WGD had occurred at that node or up to two nodes *prior* to the regime shift.

## RESULTS

### PTGR evolution and C-value evolution in angiosperms versus gymnosperms

The *PTGR* tree comprised 451 seed plants, with 28 species from 7 of 8 gymnosperm orders (Christenhusz et al. 2011) including Cycads, *Ginkgo*, conifers and Gnetales; and 423 species from 38 of 64 (59%) of angiosperm orders (APG IV 2016), including representatives from all three ANA grade lineages, Chloranthaceae, eumagnoliids, and a broad distribution of both monocots and eudicots (full tree in Appendix S2). Gymnosperm *PTGR*s ranged from < 1 to 19 μm h^−1^ (mean ± *SD* = 3.29 ± 4.34, median = 1.49 μm h^−1^), whereas angiosperm *PTGR*s ranged from < 5 to > 30,000 μm h^−1^ (mean ± *SD* = 1744 ± 3576 μm h^−1^, median = 587 μm h^−1^). The maximum likelihood (ML) reconstruction indicated that ancestral log_10_ *PTGR* of extant angiosperms was 2.44 μm h^−1^ (95% CI: 1.09-3.69) versus 0.215 μm h^−1^ (95% CI: −1.48−1.92) for extant gymnosperms (Appendix S2).

Model-based analyses of seed plant *PTGRs* and C-values favored OU models with separate optima for angiosperms and gymnosperms, accounting for > 99.9 % of the model weight in both analyses (Appendix S5, S6). Log_10_ *PTGR* optima were more than an order of magnitude faster in angiosperms (2.69 ± 0.048 μm h^−1^) than in gymnosperms (0.187 ± 0.123 μm h^−1^).

The C-value tree included 183 species from the *PTGR* tree for which DNA content data could be obtained. The resulting log_10_ C-value optimum for angiosperms (0.184 ± 0.051 pg) was more than a magnitude of order smaller than that of gymnosperms (1.231 ± 0.041 pg). Ancestral Log_10_ DNA content was also smaller for angiosperms than for gymnosperms, 0.29 pg (95% CI: −0.45−1.04) versus 1.10 pg (95% CI: −0.28−2.47), consistent with larger comparative analyses of DNA content (see Leitch and Leitch, 2013).

### Joint evolution of PTGR and ploidy

In model-based analyses of angiosperms using the empirical error rate, 89% of the model weight favored a separate and higher optimum for neo-polyploids (*N* = 68) than for diploids (*N* = 138) (model averaged log_10_ *PTGR* = 3.2 ± 0.23 vs. 2.8 ± 0.08 μm h^−1^; Table 2). In the sensitivity analysis, OU models with separate and faster *PTGR* optima for neo-polyploids than diploids received > 50% of model weight when the error calculated from *CV*s ranged from 0 to 25 %, but above 25% single-regime and BM models had the majority of the weight (Appendix S4). These are conservative results, since *SDs*, not *SEs*, were used to model error on the tree. The gymnosperm-only analysis was not performed due to low sample size (2 of 23 species were polyploid).

A survey of intraspecific cytotypes found autopolyploids had slower *PTGR* than diploids in 9 of 11 pairs and no difference in the remaining two (Binomial test, *P* = 0.002; Appendix S7b). In the nearest-relative comparisons, within-genus polyploids had slower *PTGR* than diploids in four pairs, faster *PTGR* in five, and no difference in one (Two-tailed binomial test, *P* = 0.623)(Appendix S7a).

The historical effect of number of ancient genome duplications on *PTGR* was non-significant, whether or not recent (within-genus) WGDs were included (kappa model weight > 99.9%, *N* = 451; *P >* 0.3 in both analyses).

### Joint evolution of PTGR, DNA content and ploidy

For seed plants, ordinary least squares (*OLS*) regression showed a significant negative correlation between DNA content and *PTGR* (*N =* 183, *P*<0.0001), but that result was clearly driven by the large DNA contents and slow *PTGRs* of gymnosperms relative to angiosperms (Fig. 2), because the *PGLS* regression was non-significant (Table 1). Taking these two clades separately, DNA content was negatively correlated with *PTGR* in gymnosperms in the *PGLS* regression (*N* = 23; model-averaged slope: −1.09 ± 0.49 log_10_ *PTGR*; Table 1). In angiosperms, a positive correlation using *OLS* (*N* = 161; *P*=0.0005), was non-significant using *PGLS* (Table 1). The patterns of *PTGR* and C-value evolution in seed plants can be visualized in Figure 3. In a smaller phylogenetic ANCOVA analysis, after controlling for C-value, the effect of ploidy on *PTGR* was non-significant in angiosperms (*N =* 100) and seed plants (*N =* 118) (non-significant ploidy x C-value interaction removed; Appendix S8).

**Table 1:** Phylogenetic generalized least squares regression of log_10_ *PTGR* as a function of log_10_ C-value. Only models contributing more than 1% of total weight are included. *P* values are for the slope of the regression. Gymnosperm averaged model: Log_10_ *PTGR* = 1.46 (± 0.62) −1.09 (± 0.49)*(log_10_ C-value).

| Seed plants (*N* = 183) |  |  | Angiosperms (*N* = 161) |  |  | Gymnosperms (*N* = 23) |  |  |
| --- | --- | --- | --- | --- | --- | --- | --- | --- |
| Model | Weight | *P* | Model | Weight | *P* | Model | Weight | *P* |
| kappa | 0.999 | 0.463 | kappa | 0.975 | 0.284 | OU | 0.265 | 0.020 |
|  |  |  | lambda | 0.024 | 0.221 | delta | 0.257 | 0.006 |
|  |  |  |  |  |  | kappa | 0.193 | 0.445 |
|  |  |  |  |  |  | BM | 0.121 | 0.001 |
|  |  |  |  |  |  | lambda | 0.119 | 0.327 |
|  |  |  |  |  |  | EB | 0.044 | 0.001 |

**Table 2:** Parameter estimates for angiosperm *PTGR* analyses under different evolutionary models. Note that OU1 is a single optimum model, and the rest specify separate “diploid” (paleo-polyploid) and neo-polyploid optima. BM1 and BMS models contributed <1% model weight and were excluded.

| **Model** | **$\Delta AICc$** | **Model weight** | **Diploid $\sigma^2$** | **Polyploid $\sigma^2$** | **Diploid $\alpha$** | **Polyploid $\alpha$** | **Diploid optimum** | **Polyploid optimum** |
| --- | --- | --- | --- | --- | --- | --- | --- | --- |
| **OUMA** | [332.76] | 0.373 | 0.099 |  | 0.077 | 0.074 | 2.776 | 3.285 |
| **OUMV** | 0.23 | 0.333 | 0.099 | 0.057 | 0.077 |  | 2.768 | 3.262 |
| **OUM** | 1.53 | 0.174 | 0.090 |  | 0.080 |  | 2.760 | 3.263 |
| **OU1** | 2.26 | 0.120 | 0.089 |  | 0.077 |  | 2.812 |  |
| **AVERAGED MODEL** |  | ~1.0 | 0.096 ± 0.177 | 0.082 ± 0.220 | 0.078 ± 0.213 | 0.077 ± 0.217 | 2.775 ± 0.079 | 3.217 ± 0.231 |

**Figure 2:**
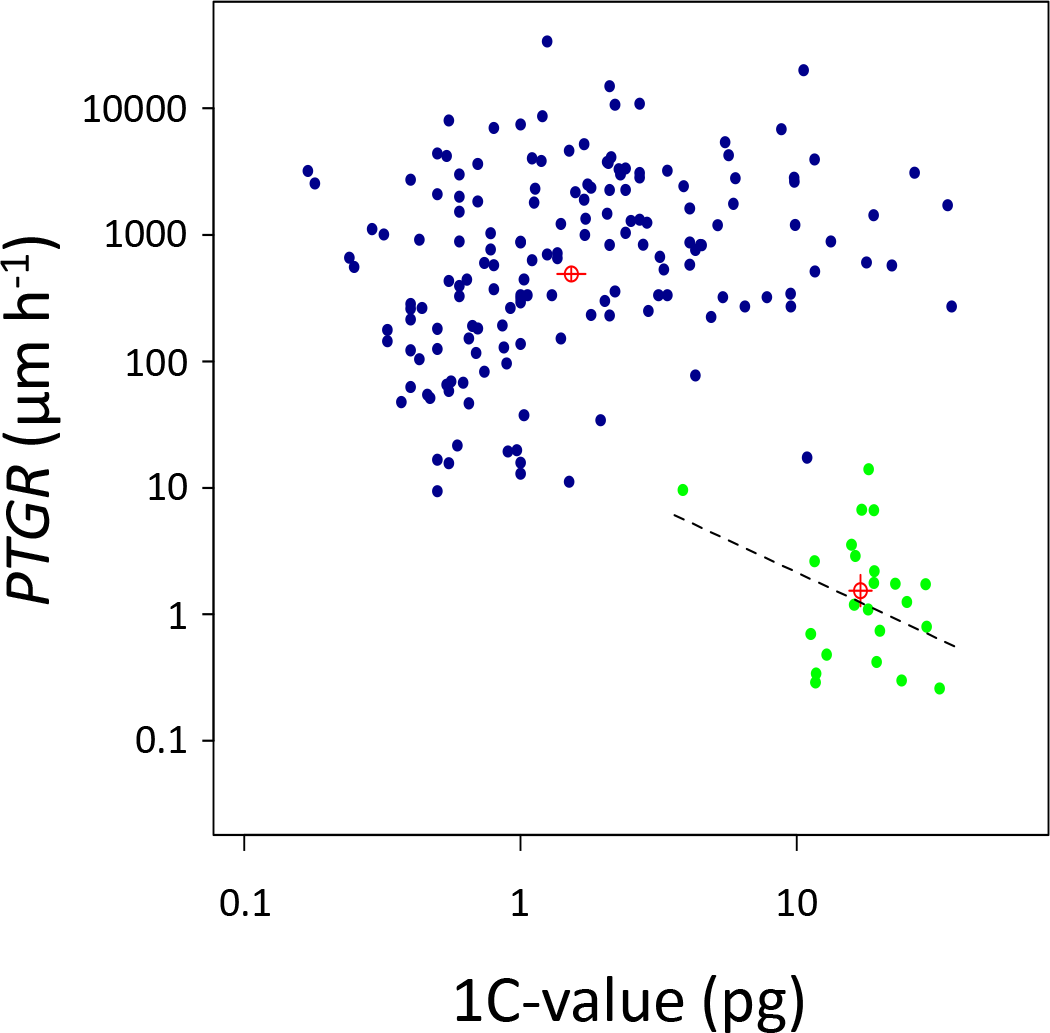
Relationship between pollen tube growth rate (*PTGR*) and DNA content (1C-value) in seed plants. The model-averaged slope of the PGLS regression is shown for gymnosperms (green points, *N* = 161), whereas slopes for seed plants (all points, *N* = 183) and angiosperms (purple points, *N* = 23) were non-significant. Optima (with standard error bars) for each group (from model-based analyses in Tables S3, S4) are included for illustrative purposes.

**Figure 3:**
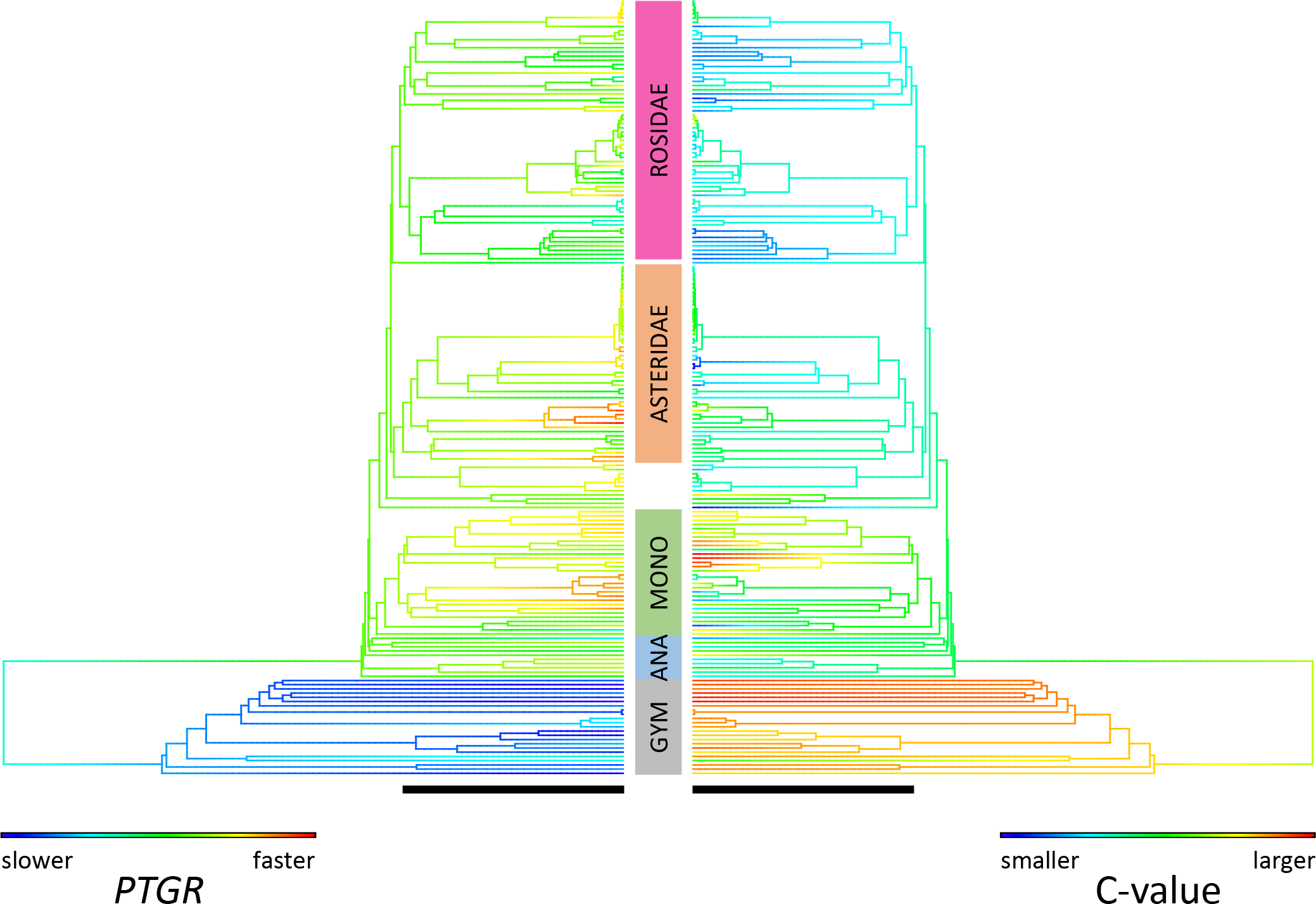
Inferred pattern of pollen tube growth rate (*PTGR*) and genome size changes in seed plants. Contour plot comparing *PTGR* evolution (left, μm h^−1^) and C-value evolution (right, picograms) (*N* = 183). Scale bar at the bottom of each phylogeny indicates 100 million years. GYM = gymnosperms; ANA = Amborellales, Nymphaeales, Austrobaileyales, Chloranthales, eumagnoliids; MONO = monocots.

### Coincident regime shifts in PTGR and DNA content

Maximum likelihood analysis of convergent evolution of *PTGRs* detected 13 distinct optima (*N* = 451 taxon tree), with 51 shifts (22 to faster and 29 to slower optima). For C-value, there were 9 distinct optima (*N* = 184 taxon tree), with 4 shifts to larger and 7 shifts to smaller optima. Regime shifts in both traits were coincident at only two nodes: a *PTGR* acceleration (from θ = 0.147 to θ = 2.47 log_10_ μm h^−1^) and genome downsizing (θ = 2.71 to θ = 0.702 log_10_ pg) in the CA of extant angiosperms; and a *PTGR* slowdown (θ = 2.78 to θ = 2.47 log_10_ μm h^−1^) and genome size decrease (θ = 0.209 to θ = −0.386 log_10_ pg) in the CA of rosids and Saxifragales (i.e. superrosids; Fig. 4). When the search was relaxed to include adjacent nodes, an additional coincidence occurred, with shift to higher *PTGR* followed by a shift to higher C-value near the base of monocots. Ancient WGDs coincided with the shifts in *PTGR* and C-value at the CA of angiosperms (above) and with a decrease in C-value in the CA of eudicots.

**Figure 4:**
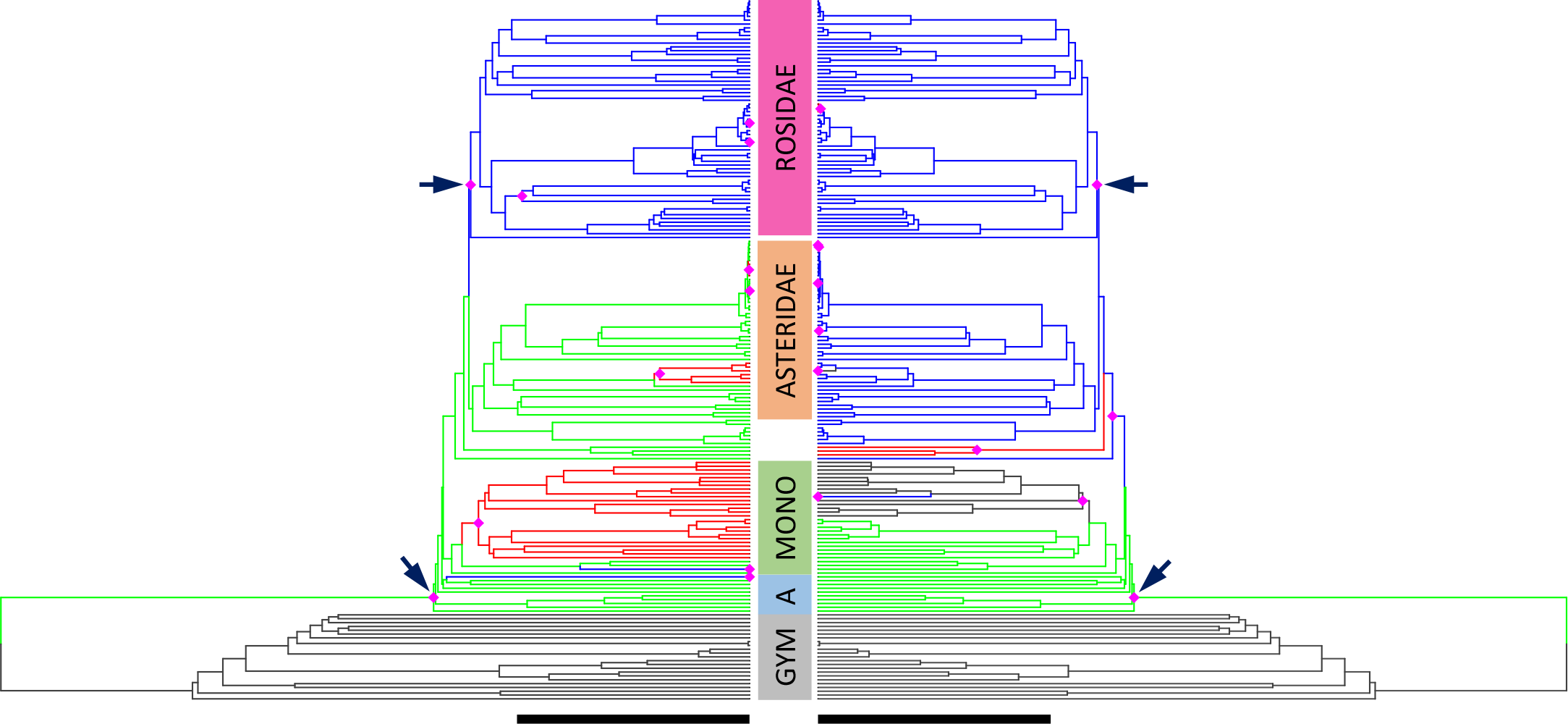
Coincident evolution of pollen tube growth rate (*PTGR*) and DNA content (C-value). Paired *SURFACE* plot showing regime shifts in *PTGR* (left) versus DNA content (right) (*N* = 183). Nodes which have experienced a regime shift along the stem leading to it are marked with magenta diamonds (not all *PTGR* shifts are shown, since *PTGR* tree has been pruned to match C-value tree). Branch colors: *gray* = seed plant ancestral optimum (*PTGR* θ = 0.147; C-value θ = 2.71); *green* = ancestral optimum for angiosperms (*PTGR* θ = 2.47; C-value θ = 0.702); *red* = derived lineages following a shift to a higher optimum than previously; *blue* = derived lineages following a shift to a lower optimum than previously. Black arrows indicate instances where shifts in *PTGR* and C-value coincide. Scale bar at the bottom of each phylogeny indicates 100 million years. GYM = gymnosperms; A = Amborellales, Nymphaeales, Autrobaileyales, Chloranthales, eumagnoliids; MONO = monocots.

## DISCUSSION

The impact of genome size on *PTGR* is determined by the magnitudes of conflicting nucleotypic and genotypic effects. Such effects depend on the mechanism of genome size change. Nucleotypic effects decelerate *PTGR* and are always present irrespective of mode of genome size change, whereas large-scale genetic effects are only possible after WGD. We predicted that angiosperms and gymnosperms should have different patterns of *PTGR* evolution based on their contrasting patterns of genome size change. Gymnosperm *PTGRs* should be most susceptible to nucleotypic effects because they have evolved large genomes sizes and WGDs have been rare. In contrast, angiosperms have evolved smaller genome sizes despite recurrent WGDs and widespread present-day polyploidy. Thus, gene duplication and sorting have played a much greater role in the evolution of angiosperm *PTGRs*, allowing genotypic effects to counterbalance or overwhelm nucleotypic effects. Below we discuss our findings in light of these expected patterns.

### *The evolution of* PTGR *in angiosperms versus gymnosperms*

We found that seed plant *PTGRs* best fit an OU model, indicating less *PTGR* variation among lineages than expected under a Brownian motion evolutionary model, with a faster optimum for angiosperms than for gymnosperms. Phylogenetic half-lives were similar (5.6 and 5.7 MY, respectively) and very short (only 3.9% and 2.3 % of their respective crown ages), indicating a strong attraction to their optimum values. Such a pattern is consistent with stabilizing selection on *PTGR* imposed by slower evolution of linked sporophytic traits, such as the timing of stigma receptivity relative to egg receptivity, pollen tube pathway length, or maternal provisioning. Gymnosperm *PTGRs* may have been constrained by a hard boundary such as by biophysical or physiological limitations, or a soft boundary, such as by lack of selection for fast rates. Angiosperms have clearly not been bound by those same limitations, given their much higher *PTGR* optimum, the convergent evolution of extremely fast *PTGRs* in many unrelated derived lineages of monocots and eudicots, and occasionally large within-genus differences in *PTGR*.

Our results suggest that most of the accelerations of angiosperm *PTGR*, and their higher *PTGR* variance relative to gymnosperms, have largely evolved *after* the origin of angiosperms and their novel pollen tube cell biology. First, estimates of angiosperm ancestral *PTGR* and ancestral optimum under OU (275 and 295 μm h^−1^, respectively) are slower than the angiosperm-wide OU optimum of 490 μm h^−1^ and the angiosperm median of 587 μm h^−1^. Secondly, the higher among-lineage variance is due to many transitions to both faster and slower *PTGR* optima within extant angiosperms. Transitions to slower rates within angiosperms are concentrated on lineages that have evolved delayed fertilization, such Fagales, orchids and others, or high selfing rates, which suggests relaxation of directional selection on *PTGR* (Williams and Reese, 2019). In contrast, gymnosperm *PTGRs* were likely ancestrally slow (Figure 4).

There are several non-mutually exclusive hypotheses for what triggered the evolution of fast *PTGRs* in angiosperms. First, Mulcahy (1979) invoked a shift to much higher intensity of pollen competition in angiosperms as a driver of the origin and continued evolution of faster growth rates. Notably, no other type of tip-growing cell in land plants (whether gametophytic or sporophytic) has evolved comparably fast tip-growth rates and none of those cell types, including gymnosperm pollen tubes, experience intense competition for resources (Williams et al., 2016). Secondly, gymnosperm *PTGRs* may be slow because they lack novel biophysical or physiological attributes of pollen tubes and/or those attributes enabled faster *PTGRs* to evolve in angiosperms (Hoekstra, 1983; Derksen et al., 1999; Fernando et al., 2005; Williams, 2008, 2009). Thirdly, with or without pollen competition, rapid *PTGRs* may have been necessary as angiosperm sporophytes transitioned to a much faster reproductive cycle (Stebbins, 1974; Williams, 2012; Williams and Reese, 2019). Finally, our results suggest a new possibility, that strong differences in genome-level processes have impacted the evolution of angiosperm *PTGRs* relative to their living and extinct seed plant relatives.

### Mechanisms of genome size change and PTGR evolution within seed plants

A major finding of this study is that angiosperm neo-polyploids evolved around a much faster *PTGR* optimum (1648 μm h^−1^) than diploids (595 μm h^−1^), despite several sources of variation in the data. First, neo-polyploids were by definition derived within genera, and their smaller sample size and shorter branch lengths reduced the power to estimate parameters relative to diploids, as reflected by the larger standard error around the neo-polyploid optimum. Nevertheless, the proportion of neo-polyploids in our data set (33% of angiosperms) is almost exactly that found in the full Wood et al. (2009) data set and similar to that in other studies (Mayrose et al., 2011; Barker et al., 2016; Landis et al., 2018).

There was also biological variability in our dataset. In our taxon sampling, we were agnostic to variation in mating systems and modes of polyploid origins, since our interest was in how *PTGR* has evolved in natural stabilized polyploids. In retrospect, our sample does seem representative. Of 14 angiosperm polyploids whose mode of origin has been studied, seven were autopolyploid and seven allopolyploid, similar to the nearly-equal proportions found by Barker et al., 2016. Furthermore, among 16 polyploids for which mating system has been studied, eight were fully outcrossing, seven were self-compatible (two autogamous, two mixed mating, and three unknown), and one was apomictic – a not unusual distribution (Goodwillie et al., 2005; Gibbs, 2014; Ashman et al., 2014). Thus, our taxon sampling seems not be have been greatly biased. Even with such information, predicting the magnitude of genetic variation in polyploids is not so simple. For example, autotetraploids originate with a subset of the genetic variation in the diploid progenitor population but they often outcross and hybridize, whereas allopolyploids can be highly heterozygous when they originate, but often are highly selfing (Stebbins, 1974; Soltis and Soltis, 1999; Barringer 2007; Whitney et al., 2010). Hence, despite several sources of heterogeneity, the faster *PTGR* optimum of neo-polyploids indicates that *PTGR* acceleration evolves either at the time of WGDs or during the time period in which the descendant species retain a polyploid chromosome number.

The closest approximation of the initial effect of polyploidy on *PTGR* is the comparison of diploids with their intraspecific, autopolyploid cytotypes. In all 11 pairs, *PTGRs* of autopolyploid cytotypes were slower than or equal to those of their intraspecific diploid progenitors. We should re-emphasize that all studies involved in vivo crosses among diploid sporophytes (1x pollen on 2x pistils) compared to crosses among tetraploid sporophytes (2x pollen on 4x pistils), in keeping with our goal of generalizing effects on *PTGR* in stabilized polyploids. Nucleotypic effects acting to slow *PTGR* should be most apparent in autopolyploids at inception, because there is lower potential for heterosis. Thus, the lack of any examples of faster *PTGR* in neo-autotetraploid cytotypes than in their diploid progenitors suggests that increased gene dosage by itself generally does not initially fully offset nucleotypic effects.

Nucleotypic effects on *PTGR* could be substantial. Tube size affects *PTGR* in a linear fashion, because larger tubes must make more tube wall per unit time, and since tube diameter is constant during growth, the rate of wall production is directly proportional to tip extension rate (Williams et al., 2016). Kostoff & Prokofieva (1935) reported in vivo pollen tubes to be 39% larger in diameter in an allotetraploid *Nicotiana* relative to the mean of its presumed diploid progenitors, and Iyengar (1938) found 8-53% larger tube diameters in tetraploid versus diploid species of *Gossypium*.

Taken together our results suggest that nucleotypic effects are strong and act as a brake on *PTGR* at inception (intraspecific polyploid analysis), but as neo-polyploids become stabilized and persist over time, nucleotypic effects are more than offset by genotypic effects (within-genus pairs and model-based analyses) which often produce faster *PTGRs* in angiosperms.

We found that DNA content has evolved around a significantly lower optimum in angiosperms than in gymnosperms, even though angiosperms have a broad range of DNA C-values that encompass the entire range of seed plant genome sizes (Fig. 3; see Leitch and Leitch, 2013 for a larger survey). Angiosperms also have great variation in ploidy level, a history of speciation by polyploidy, and much evidence of past genome duplication (Ahuja, 2005; Wood et al., 2009; Husband et al., 2013; Van de Peer et al., 2017; Landis et al., 2018). There were at least 1-7 WGDs in the lineages leading from the seed plant root to each of the tips in our *PTGR* tree, and 33% of taxa (68/206 angiosperms versus 2/23 gymnosperms) were identified as neo-polyploids. The often low DNA content and high ploidy levels of angiosperms are not surprising given that genome duplication is commonly followed by rapid loss of DNA sequences, gene fractionation by large-scale deletions, biased retention of genes with beneficial dosage effects, and ultimately a return to an apparent diploid state in sporophytes (Conant and Wolfe, 2008; Conant et al., 2014; Freeling et al., 2015; Dodsworth et al., 2016; Wendel et al., 2018). Thus, one explanation for the much faster *PTGRs* of angiosperms relative to gymnosperms is that widespread gene duplication by WGDs have often enabled transgressive evolution of faster *PTGRs* leading to the observed pattern of convergent evolution of extremely fast *PTGRs* in many unrelated lineages of monocots and eudicots.

WGDs have been rare in gymnosperms (Ahuja, 2005; Leitch et al., 2005; Wood et al., 2009; Soltis et al., 2009; Husband et al., 2013; Leitch and Leitch, 2013; Lee and Kim, 2014) and their high DNA contents are thought to be due mainly to high transposon activity without repeated rounds of genome duplication (Leitch & Leitch, 2013; Lee and Kim, 2014). Hence, gymnosperms may have experienced the nucleotypic effects of higher DNA content on pollen tube dimensions, which is predicted to reduce *PTGR*, without the potential for counter-balancing effects, such as initially higher gene dosage and heterozygosity followed by gene sorting during the diploidization process. Our finding of a negative correlation between *PTGR* and DNA content in gymnosperms, but not in angiosperms supports that hypothesis.

Though gymnosperm *PTGRs* are likely affected by tube sizes, nucleotypic effects do not account for the magnitude of the difference in their slow *PTGRs* relative to those of angiosperms. Gymnosperm pollen tubes can range up to 300 μm in diameter (Coulter and Chamberlain, 1928; Gifford and Foster, 1989), but many species of siphonogamous conifers and Gnetales have angiosperm-like pollen tube diameters in the 10 to 20 μm range. Yet no gymnosperm has evolved a *PTGR* faster than 20 µm h^−1^. It has been argued that their pecto-cellulosic wall structure is a limitation relative to angiosperm pollen tube walls, which use the plasma membrane-bound enzymes callose synthase and pectin-methylesterase in a novel way to more rapidly synthesize a strong and durable tube cell wall and callose plugs (Derksen, 1999, Abercrombie et al., 2012; Wallace and Williams, 2017). However, other types of pecto-cellosic tip-growing cells, such as root hairs, grow faster than gymnosperm pollen tubes (Williams et al., 2016). Thus, it seems likely that the extremely slow growth rates of gymnosperm pollen tubes reflect an ancestrally antagonistic relationship between maternal tissues and pollen tubes that functioned as invasively growing rhizoids, coupled with a lack of selection for faster growth rate due to the absence of pollen competition and a long period between pollination and fertilization. Our results also suggest a lack of opportunity for genotypic effects to evolve due to the rarity of WGDs.

### Conclusions

Studies across the tree of life have consistently shown that ploidy level and DNA content are correlated with cell size and metabolic rate (Cavalier-Smith, 1978; Gregory, 2001; Cavalier-Smith, 2005). Pollen tube dimensions and energetics affect the amount of cell wall material produced per unit of growth and the rate at which cell wall is produced, which together determine *PTGR*. In gymnosperms, *PTGR* was negatively correlated with genome size, but in angiosperms, where the effects of WGDs are much more prevalent, there was no such correlation, and neo-polyploids evolved around a higher *PTGR* optimum than diploids. These results support the expectation that genome size increases incur nucleotypic effects that act as a brake on growth rate. The degree to which genotypic effects counterbalance nucleotypic effects depends on the historical nature and time since genome size increase in any particular lineage. Understanding causal relationships between genome size, ploidy and *PTGR* will involve mechanistic studies of tube cell dimensions and wall synthesis rates in haploid and polyploid gametophytes. On the other hand, there appears to be great variation in the tug of war between genotypic and nucleotypic effects, and there are likely to be deeper evolutionary patterns underlying that variation.

## ACKNOWLEDGEMENTS

We thank B. O’Meara and J. Beaulieu for advice on phylogenetic analyses, I. Leitch for data on DNA content, and J. Edwards and M. Rankin for assistance in the lab. We are tremendously grateful to several anonymous reviewers for their perceptive and useful comments. Partial support to J.B.R. was provided by National Science Foundation award IOS 1052291 to J.H.W.

## Authors Contributions

J.B.R. and J.H.W. jointly conceived of the study and wrote the paper;J.B.R. collected data on genome sizes and ploidy levels, constructed the phylogenetic tree and performed all comparative analyses; J.H.W. collected data on *PTGRs* and diploid-autopolyploid *PTGRs*.

## Data Accessibility Statement

Scripts written during the creation of this manuscript are available on GitHub: https://github.com/jbr1848/PTGR.genome.evolution. The phylogenetic tree created during this study can be found on TreeBase: http://purl.org/phylo/treebase/phylows/study/TB2:S24291.

## Additional Supporting Information may be found online in the supporting information section at the end of the article

**Appendix S1:**
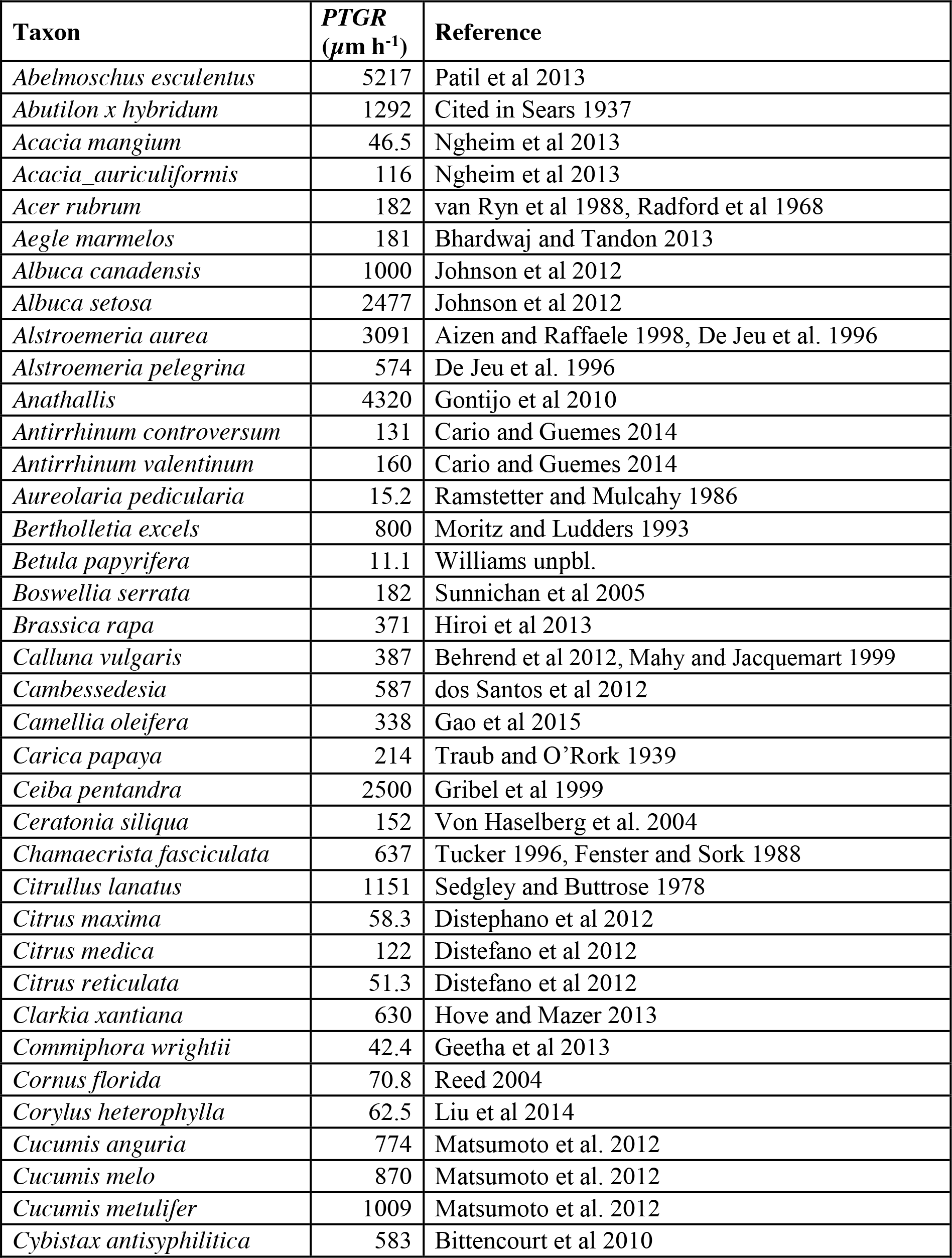
119 additional *PTGR* values and references not reported in Williams 2012.

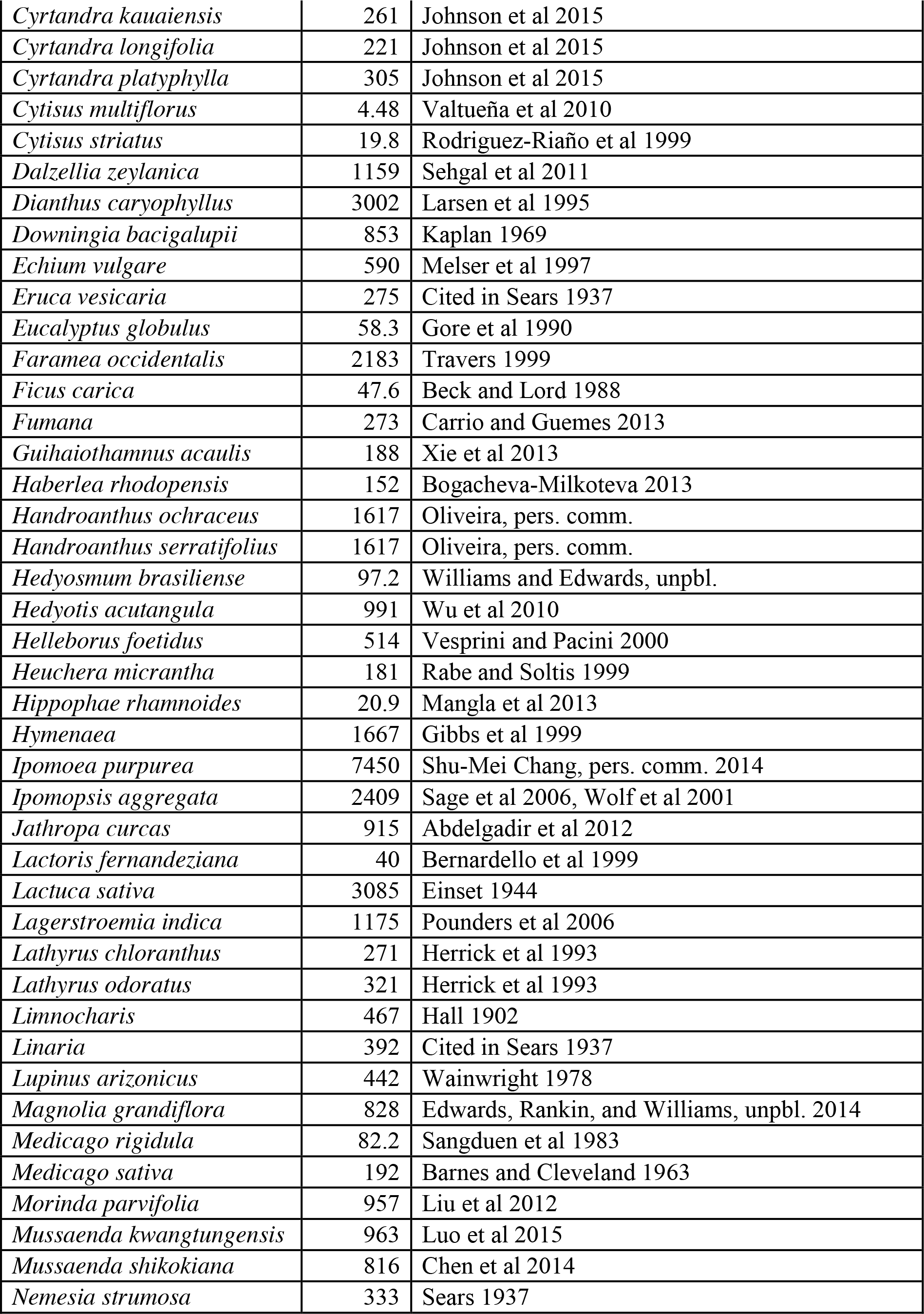

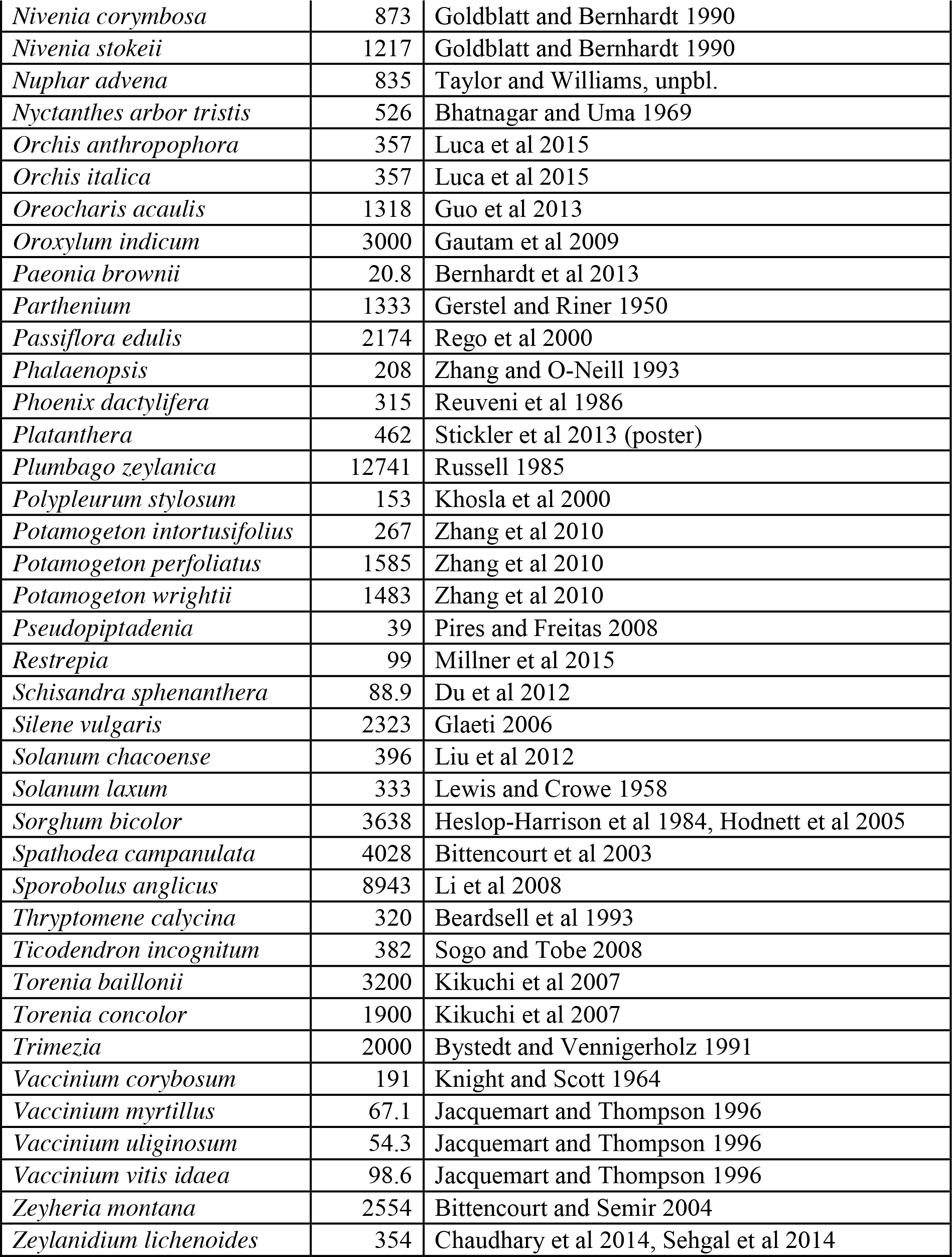

**Appendix S2:**
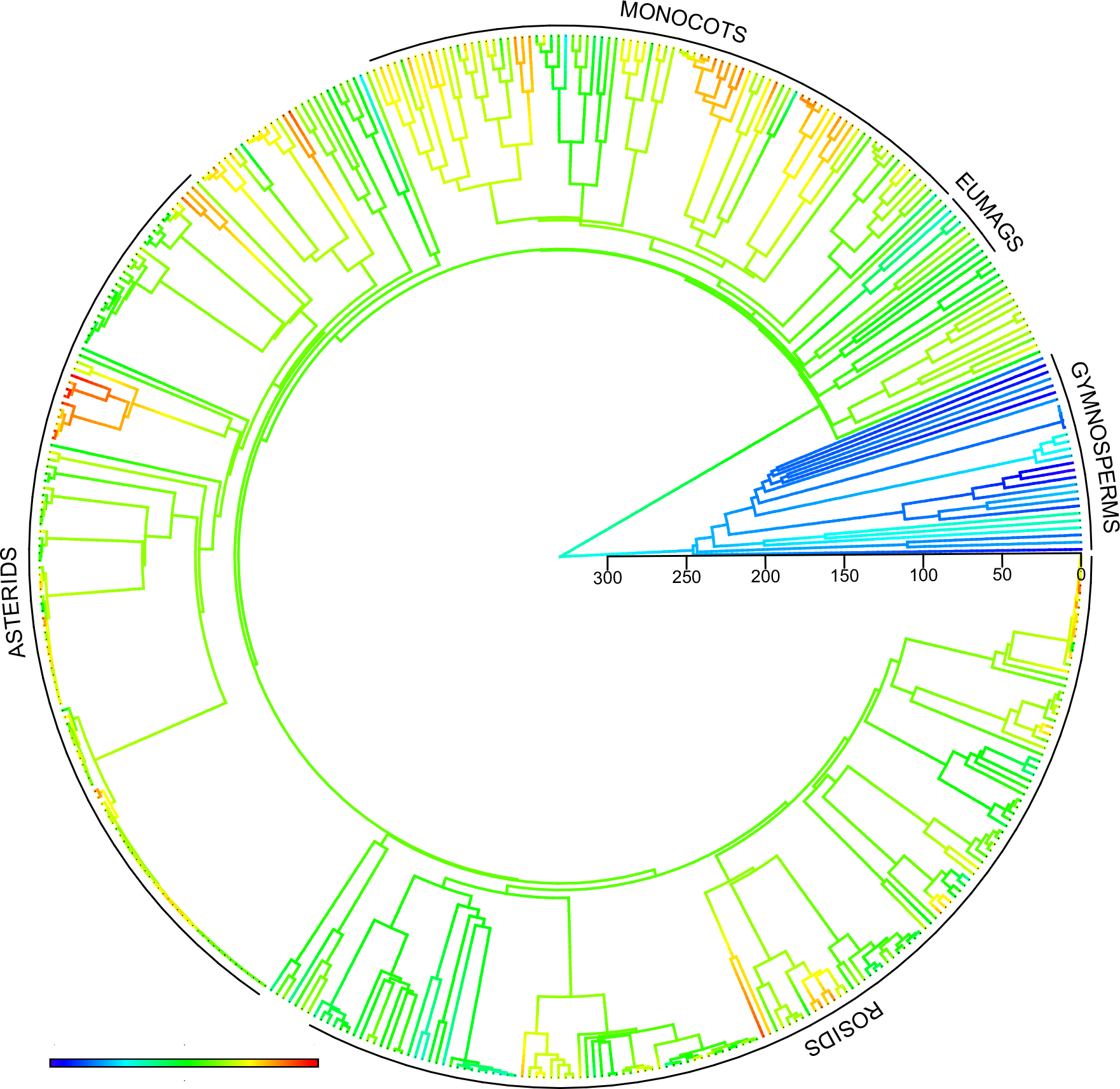
Pollen tube growth rate (*PTGR*) evolution across Spermatophytes. Contour plot showing reconstructed history of *PTGR*. Cool colors indicate *PTGR*s closer to the minimum value in seed plants while warm colors indicate *PTGR*s closer to the maximum value in seed plants. Scale bar indicates millions of years before present.

**Appendix S3:** Summary statistics for pollen tube growth rate (*PTGR*) of *Magnolia grandiflora*.

| **Statistic** | ***PTGR*** |
| --- | --- |
| raw mean ($N = 25$ crosses) | 827.6 $\mu\text{m h}^{-1}$ |
| raw SD ($N = 25$) | 141.3 $\mu\text{m h}^{-1}$ |
| raw CV | 0.1708 |
| Log(10) mean | 2.912 $\mu\text{m h}^{-1}$ |
| transformed SD | 0.0689 $\mu\text{m h}^{-1}$ |
| transformed CV | 0.0237 |

**Appendix S4:** Sensitivity analysis for the magnitude of log_10_ *PTGR* error estimates. Values in each column represent model weights from separate analyses of angiosperm diploids (*N* = 138) vs. polyploids (*N* = 68). Column headings indicate the coefficient of variation (CV), ranging from zero to 0.50, used to calculate estimated species-specific standard deviations around *PTGRs* in each analysis. The best-fitting model at each CV is indicated in bold. ^a^, Empirically-determined CV of *Magnolia grandiflora*.

| Model | Coefficient of Variation |  |  |  |  |  |
| --- | --- | --- | --- | --- | --- | --- |
|  | 0.00 | 0.0237<sup>a</sup> | 0.05 | 0.10 | 0.25 | 0.50 |
| OUMV | **0.376** | **0.373** | **0.459** | **0.461** | 0.119 | 0.045 |
| OUMA | 0.363 | 0.333 | 0.307 | N/A | 0.172 | 0.070 |
| OUM | 0.157 | 0.174 | 0.137 | 0.309 | 0.212 | 0.080 |
| OU1 | 0.104 | 0.120 | 0.097 | 0.230 | **0.390** | 0.188 |
| BM1 | 1.74E-31 | 1.77E-25 | 3.20E-20 | 4.58E-14 | 0.089 | **0.528** |
| BMS | 5.45E-32 | 5.52E-26 | 4.81E-24 | 1.30E-12 | 0.017 | 0.089 |

**Appendix S5:** *PTGR* evolution in gymnosperms vs. angiosperms. Selective regime 1 represents gymnosperms (*N =* 28) and selective regime 2 represents angiosperms (*N =* 423). Models representing <1% of the model weight are excluded.

| model | $\Delta AIC_c$ | model weight | sigma sq1 | alpha1 | sigma sq2 | alpha2 | optimum1 | se1 | optimum2 | se2 |
| --- | --- | --- | --- | --- | --- | --- | --- | --- | --- | --- |
| **OUMV** | (822.1) | 0.338 | 0.091 | 0.122 | 0.139 | 0.122 | 0.187 | 0.123 | 2.690 | 0.048 |
| **OUM** | 0.040 | 0.332 | 0.137 | 0.124 | 0.137 | 0.124 | 0.188 | 0.150 | 2.690 | 0.047 |
| **OUMA** | 0.051 | 0.330 | 0.091 | 0.120 | 0.091 | 0.123 | 0.187 | 0.123 | 2.690 | 0.048 |
| **AVERAGED MODEL** |  | ~1 | 0.106 | 0.122 | 0.122 | 0.123 | 0.187 | 0.132 | 2.690 | 0.048 |

**Appendix S6:**
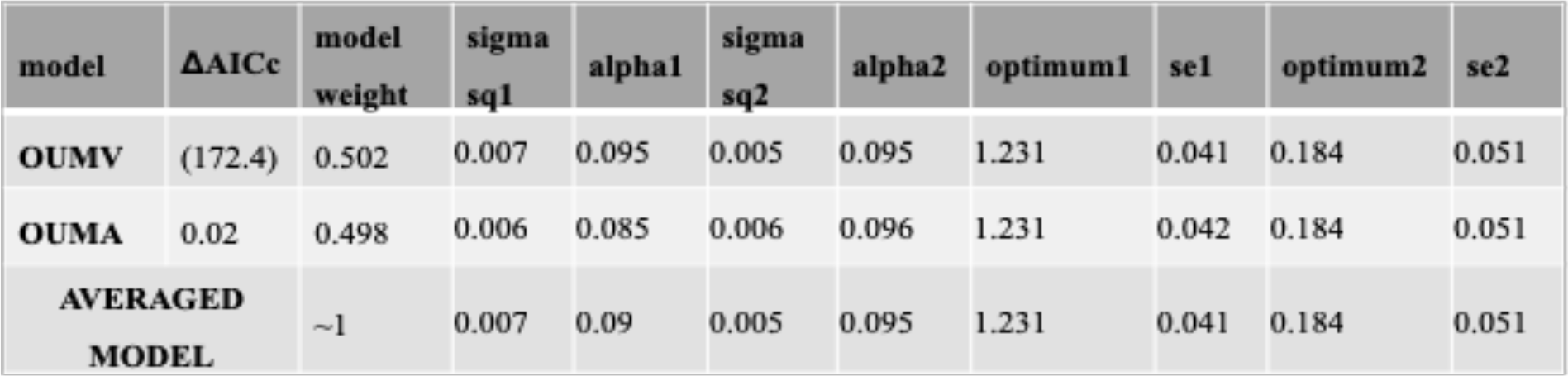
C-value evolution in gymnosperms vs. angiosperms. Selective regime 1 represents gymnosperms (*N* = 23) and selective regime 2 represents angiosperms (*N* = 161). Models representing <1% of the model weight are excluded.

**Appendix S7: Closely-related taxon analyses**

**Appendix S7a.** Closely-related species pairs extracted from ploidy dataset. *PTGRs* in µm h^−1^. Binomial test (*P* = 0.623; *N* = 10).

| DIPLOID<br>TAXON | POLYPLOID<br>TAXON | DIPLOID<br>*PTGR* | POLYPLOID<br>*PTGR* | FASTER<br>TAXON |
| --- | --- | --- | --- | --- |
| *Anagallis arvensis* | *Anagallis monelli* | 233.33 | 105.56 | diploid |
| *Hemerocallis thunbergii* | *Hemerocallis fulva* | 4166.67 | 6266.67 | polyploid |
| *Ipomoea purpurea* | *Ipomoea batatas* | 7450 | 4625 | diploid |
| *Iris mandshurica* | *Iris pseudacorus* | 278.65 | 4255.50 | polyploid |
| *Lythrum junceum* | *Lythrum salicaria* | 722.22 | 493.60 | diploid |
| *Medicago rigidula* | *Medicago sativa* | 82.23 | 192.17 | polyploid |
| *Prunus avium* | *Prunus domestica* | 260.88 | 177.5 | diploid |
| *Tabebuia rosea* | *Tabebuia chrysotricha* | 1111.11 | 1342.45 | polyploid |
| *Trifolium pratense* | *Trifolium polymorphum* | 103.89 | 444.44 | polyploid |
| *Ulmus pumila* | *Ulmus americana* | 56.25 | 56.25 | equivocal |

**Appendix S7b.** Intraspecific diploid-polyploid cytotypes taken from the literature. All are autopolyploids. Binomial test, *N* = 11, *P* = 0.0020. Percent difference is calculated relative to the diploid.

| REF. | TAXON | DIPLOID<br>*PTGR* | POLYPLOID<br>*PTGR* | POLYPLOID<br>+/- (% DIFF) |
| --- | --- | --- | --- | --- |
| 1 | *Beta vulgaris*<br>2x,4x | 241.2 $\mu\text{m/h}$ | 142.7 $\mu\text{m/h}$ | slower (-69%) |
| 2 | *Cucumis melo*<br>2x,4x | “no difference” |  | equivocal |
| 3 | *Datura stramonium* 2x,4x | 2953.7 $\mu\text{m/h}$ | 2812.5 $\mu\text{m/h}$ | slower (-4.8%) |
| 4 | *Lactuca sativa*<br>2x,4x | “faster” | “slower” | slower |
| 5 | *Malus domestica*<br>2x,4x | 3.8 units/96 h | 3.1 units/96 h | slower (-18.4%) |
| 6 | *Malus domestica*<br>2x,3x | 682 $\mu\text{m/h}$ | 465 $\mu\text{m/h}$ | slower (-31.8%) |
| 7 | *Secale cereale*<br>2x,4x | 12.24 units/h | 12.08 units/h | slower (-1.3%) |
| 8 | *Solanum* sp. 2x,4x | “faster” | “slower” | slower |
| 9 | *Trifolium pratense*<br>2x,4x | 2322 $\mu\text{m/h}$ | 1950 $\mu\text{m/h}$ | slower (-16%) |
| 10,11 | *Zea mays* 2x,4x | Slower pollen germination and<br>pollen tube growth rate in 4x |  | slower |

**Appendix S8: Phylogenetic ANCOVA results.** Models comprising <1% of the model weight are excluded.

**Appendix S8a:** Angiosperms only (*N* = 100).

| **Full model** |  |  |  |  |
| --- | --- | --- | --- | --- |
| **<u>Model</u>** | **<u>Weight</u>** | **<u>P<sub>C-value</sub></u>** | **<u>P<sub>ploidy</sub></u>** | **<u>P<sub>interaction</sub></u>** |
| kappa | 0.688 | 0.028 | 0.565 | 0.334 |
| OU | 0.287 | 0.005 | 0.679 | 0.414 |
| lambda | 0.025 | 0.076 | 0.867 | 0.626 |
| Model averaged slope for C-value: $0.399 \pm 0.167$ | | | | |

| **No interaction** |  |  |  |
| --- | --- | --- | --- |
| **<u>Model</u>** | **<u>Weight</u>** | **<u>P<sub>C-value</sub></u>** | **<u>P<sub>ploidy</sub></u>** |
| kappa | 0.652 | 0.046 | 0.748 |
| OU | 0.313 | 0.006 | 0.895 |
| lambda | 0.035 | 0.083 | 0.979 |
| Model averaged slope for C-value: $0.344 \pm 0.153$ | | | |

| **C-value only** |  |  |
| --- | --- | --- |
| **<u>Model</u>** | **<u>Weight</u>** | **<u>P<sub>C-value</sub></u>** |
| kappa | 0.642 | 0.045 |
| OU | 0.322 | 0.005 |
| lambda | 0.036 | 0.079 |
| Model averaged slope for C-value: $0.344 \pm 0.153$ | | |

**Appendix S8b:**
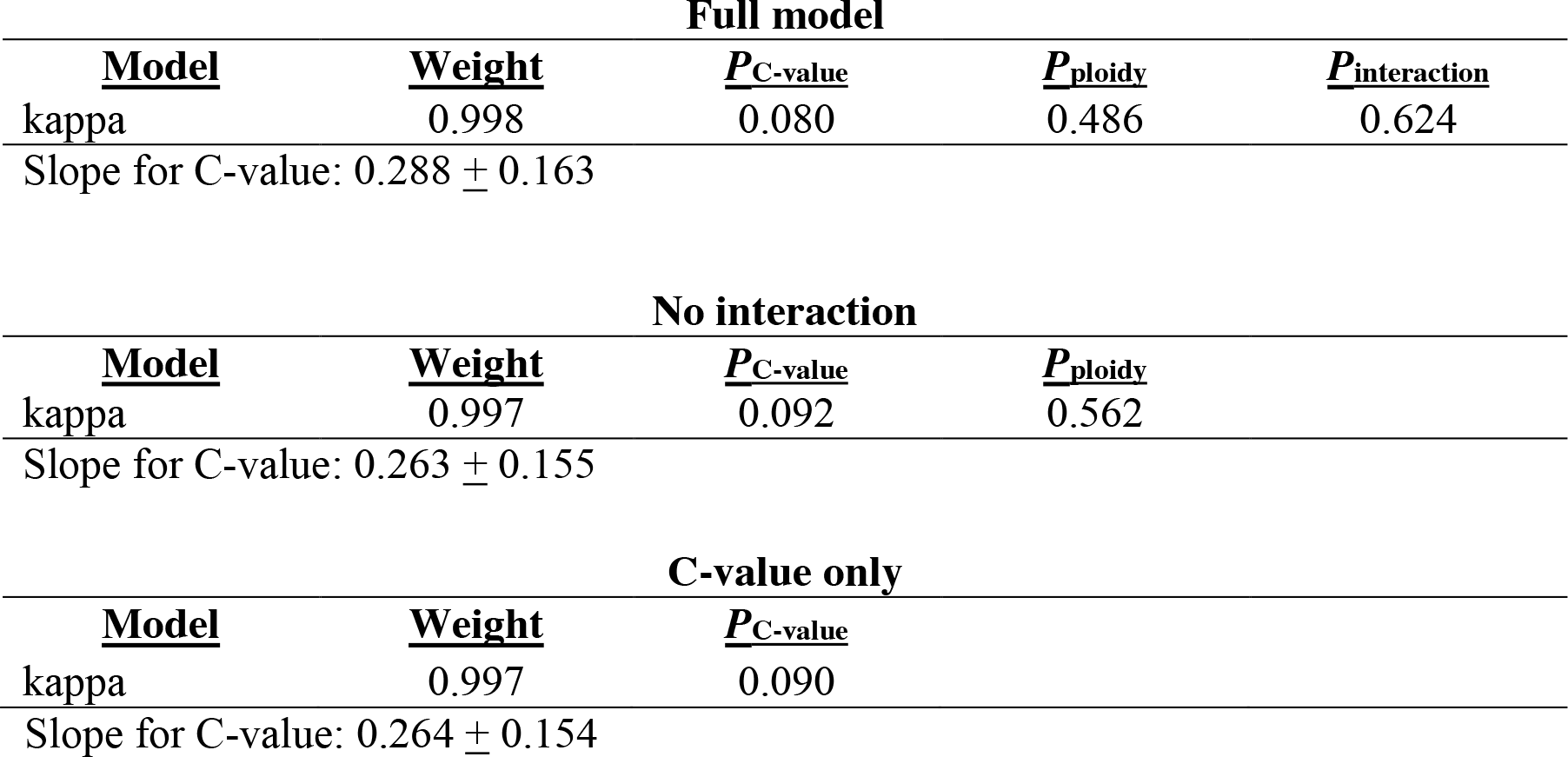
All seed plants (*N* = 118).

| **Full model** |  |  |  |  |
| --- | --- | --- | --- | --- |
| **<u>Model</u>** | **<u>Weight</u>** | **<u>P<sub>C-value</sub></u>** | **<u>P<sub>ploidy</sub></u>** | **<u>P<sub>interaction</sub></u>** |
| kappa | 0.998 | 0.080 | 0.486 | 0.624 |
| Slope for C-value: $0.288 \pm 0.163$ | | | | |

| **No interaction** |  |  |  |
| --- | --- | --- | --- |
| **<u>Model</u>** | **<u>Weight</u>** | **<u>P<sub>C-value</sub></u>** | **<u>P<sub>ploidy</sub></u>** |
| kappa | 0.997 | 0.092 | 0.562 |
| Slope for C-value: $0.263 \pm 0.155$ | | | |

| **C-value only** |  |  |
| --- | --- | --- |
| **<u>Model</u>** | **<u>Weight</u>** | **<u>P<sub>C-value</sub></u>** |
| kappa | 0.997 | 0.090 |
| Slope for C-value: $0.264 \pm 0.154$ | | |

